# Nutlin-3a induces *KRAS* mutant lung cancer specific methuosis-like cell death that is dependent on GFPT2

**DOI:** 10.1101/2023.05.25.542296

**Authors:** Dasom Kim, Dongwha Min, Joohee Kim, Min Jung Kim, Yerim Seo, Byung Hwa Jung, Seung-Hae Kwon, Ji-Yun Lee

## Abstract

Oncogenic *KRAS* mutation, the most frequent mutation in non-small cell lung cancer (NSCLC), is an aggressiveness risk factor and leads to the metabolic reprogramming of cancer cells by promoting glucose, glutamine, and fatty acid absorption and glycolysis. Lately, sotorasib was approved by the FDA as a first-in-class *KRAS*-G12C inhibitor. However, sotorasib still has a derivative barrier, which is not effective for other *KRAS* mutation types, except for G12C. Additionally, resistance to sotorasib is likely to develop, demanding the need for alternative therapeutic strategies. In this study, we found that nutlin-3a, an MDM2 antagonist, inhibited the KRAS-PI3K/Akt-mTOR pathway and disrupted the fusion of both autophagosomes and macropinosomes with lysosomes. This further elucidated non-apoptotic and catastrophic macropinocytosis associated methuosis-like cell death, which was found to be dependent on GFPT2 of the hexosamine biosynthetic pathway, specifically in *KRAS* mutant NSCLC cells. These results indicate the potential of nutlin-3a as an alternative agent for treating *KRAS* mutant NSCLC cells.

## Introduction

The incidence and mortality rates of lung cancer increases every year. Because lung cancer is difficult to detect at an early stage, it is often diagnosed at an advanced stage, as against other cancers, and its prognosis remains poor. Non-small cell lung cancer (NSCLC), the most common (∼ 85%) histological subtype of lung cancer, is characterized by various oncogenic mutations, such as those in *KRAS* (25%), *EGFR* (23%), *PIK3CA* (3%), *BRAF* (3%), *MET* (2%), and *ERBB2* (1%); these mutations play a major role in driving cancer progression (Chevallier *et al*, 2021; Yang *et al*, 2021). These driver mutations have been used as molecular targets for drug development. Among them, *KRAS* mutations are the most common mutations, with mutations in amino acid residues G12, G13, L19, D33, or Q61 (Yang *et al*, 2021; CGARN, 2014). Despite several efforts to develop drugs that target KRAS directly and indirectly, it was considered undruggable until 2021 when sotorasib (Lumakras) was approved by the U.S. FDA as a first-in-class KRAS-G12C inhibitor (AACR, 2021; Skoulidis, 2021). However, sotorasib still has a derivative barrier, which has no effectiveness for other types of *KRAS* mutations, except for G12C, and resistance is bound to develop eventually, warranting the need for other therapeutic strategies.

Recently, specific consideration has been given to the role of cellular metabolism in cancer as an alternative treatment because the reprogramming of tumor metabolism is one of the hallmarks of cancer cells (Hanahan & Weinberg, 2011; Vander *et al*, 2009). *KRAS* mutated cancer cells, including NSCLC cells, display distinct metabolic reprogramming for cancer cell growth, proliferation, and survival (Kerr & Martins *et al*, 2018; Kerk *et al*, 2021; Kimmelman *et al*, 2015) and can stimulate nutrient scavenger processes such as macropinocytosis and autophagy, which generate endogenous vesicles, macropinosomes, and autophagosome (Kimmelman & White, 2017; Recouvreux & Commisso, 2017; Kim *et al*, 2011). These vesicles eventually merge with lysosomes to break down nutrients and provide pools of nucleotides, free amino acids, lipids, and glucose to cancer cells for use in macromolecule syntheses via anabolic pathways (Recouvreux & Commisso, 2017). Active oncogenic RAS in neuroblastoma, glioblastoma, and colorectal cancer is associated with a novel type of caspase independent cell death, methuosis, which is characterized by the presence of huge cytoplasmic vacuoles derived from the abnormal growth of the macropinosome (Maltese & Overmeyer, 2014; Song *et al*, 2020; Overmeyer *et al*, 2011). Compared with normal macropinosomes that dynamically engage with cytoskeletal elements and are either tagged for recycling or fused with lysosomes, dysfunctional macropinosomes shown in methuosis fuse each other to form huge vacuoles, resulting in the rupture of cells due to swelling. Although the exact mechanism underlying this phenomenon is currently unknown, the use of methuosis has been suggested for treating *KRAS* mutant cancer (Overmeyer *et al*, 2011; Bhanot *et al*, 2010; Dendo *et al*, 2018).

In this study, we found that nutlin-3a, an MDM2 antagonist, inhibited the KRAS-PI3K/Akt-mTOR pathway and disrupted the fusion of both autophagosomes and macropinosomes with lysosomes, resulting in non-apoptotic and catastrophic micropinocytosis associated methuosis-like cell death, which was dependent on GFPT2 of the hexosamine biosynthetic pathway (HBP), in *KRAS* mutant NSCLC cells specifically; the results were further verified *in vivo*.

## Results

### Nutlin-3a suppresses *KRAS* mutant NSCLC cell viability and proliferation via the KRAS-PI3K/Akt pathway

We conducted 3-(4,5-dimethylthiazol-2-yl)-2,5 diphenyltetrazolium bromide (MTT) and colony formation assays to investigate the effect of nutlin-3a on the viability and survival of NSCLC cells and normal bronchial epithelial cells (BEAS-2B). Only *KRAS* mutant, but not wild type NSCLC cells, showed a significant decrease in viability (Fig 1A) and colony formation ability (Fig 1B) after nutlin-3a treatment. The IC_50_ values of nutlin-3a are summarized in Table EV1. To understand the effect of nutlin-3a on KRAS signaling in *KRAS* mutant NSCLC cells, KRAS activity (Fig 1C) as well as KRAS downstream signaling (Fig 1D) were examined. Nutlin-3a treatment decreased both the total KRAS and KRAS–GTP activation level as well as the levels of PI3K p110α, p-Akt, and c-MYC, but not of p-MEK and p-ERK, in *KRAS* mutant NSCLC cells

**Figure 1.**
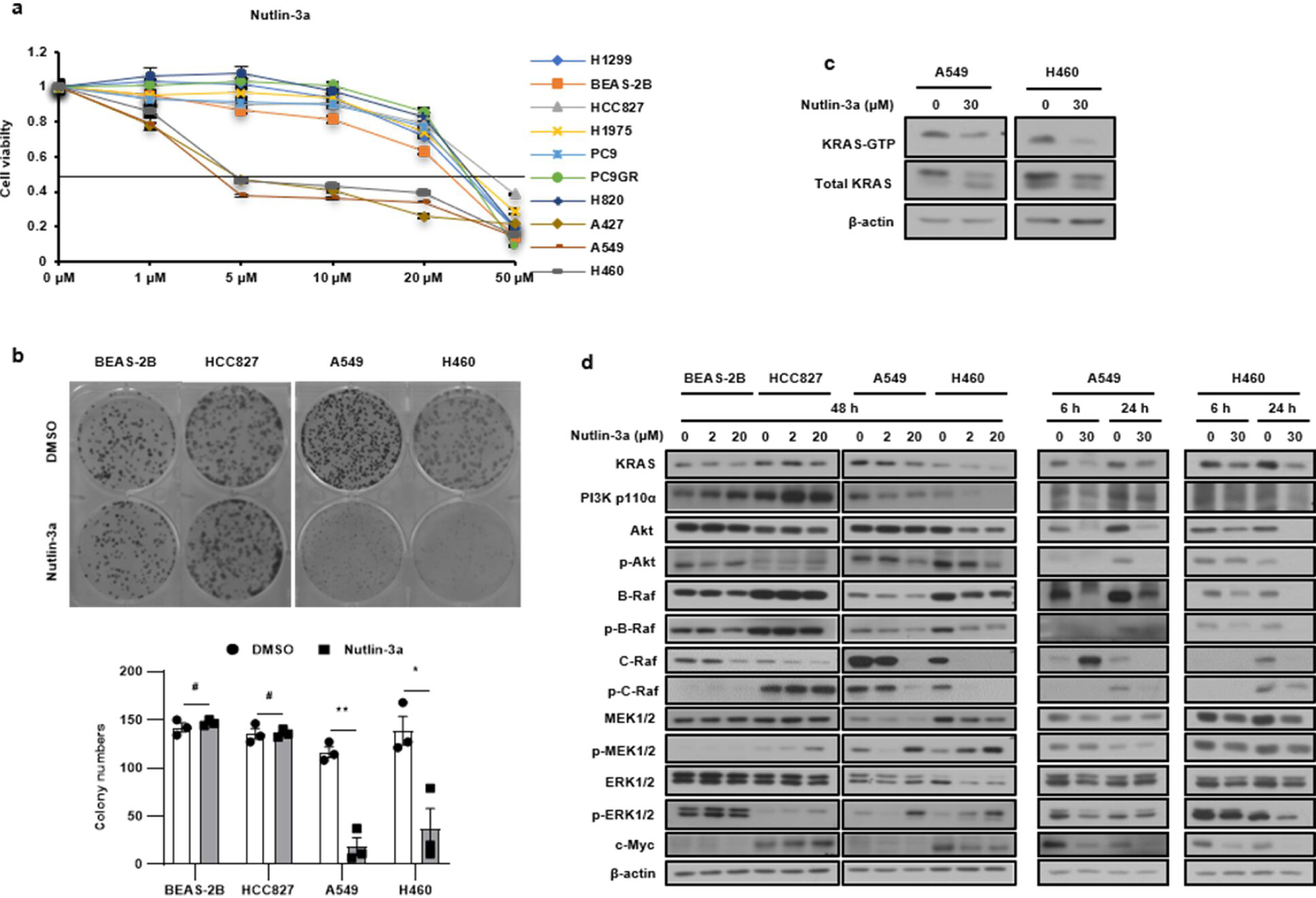
Nutlin-3a reduces *KRAS* mutant NSCLC cell viability via the KRAS-PI3K/Akt pathways. **A** Cell viability determined using MTT assay after nutlin-3a treatment for 48 h. **B** Cell proliferation examined using the colony formation assay after nutlin-3a treatment (2 μM) for 14 days (top); quantification of colony formation (bottom). **C** KRAS-GTP form detected by the KRAS activation assay after nutlin-3a treatment for 24 h. **D** Reduced expression of KRAS–PI3K/Akt and c-Myc, but not MEK/ERK, revealed by western blotting. *p<0.05, **p<0.01, ***p<0.001 compared with control.

### Nutlin-3a disrupts autophagosome-lysosome fusion

Inactivation of Akt suppresses mTOR signaling, resulting in the induction of autophagy. The mTOR pathway was examined by western blotting because the PI3K/Akt pathway was downregulated by nutlin-3a in *KRAS* mutant NSCLC cells. Nutlin-3a treatment induced autophagy by reducing p-mTOR level and downstream activity (Fig 2A). In addition, the levels of the autophagosomal markers LC3 and p62 were examined by western blotting after treatment with nutlin-3a, with or without bafilomycin A1, to investigate the effect of nutlin-3a on autophagic flux. The level of LC3-Ⅱ increased and that of p62 decreased when only nutlin-3a was administered, but co-treatment with bafilomycin A1 and nutlin-3a resulted in the accumulation of both LC3-Ⅱ and p62 in *KRAS* mutant NSCLC cells (Fig 2B). These results suggest that autophagy was induced by nutlin-3a. The mRFP-GFP-LC3 plasmid was used to further validate autophagic flux in live cells by confocal laser scanning microscopy (Fig 2C). Only the number of autophagosomes (yellow puncta) and not autolysosomes (red puncta) were increased by nutlin-3a treatment, indicating that nutlin-3a disrupts the fusion of autophagosomes and lysosomes.

**Figure 2.**
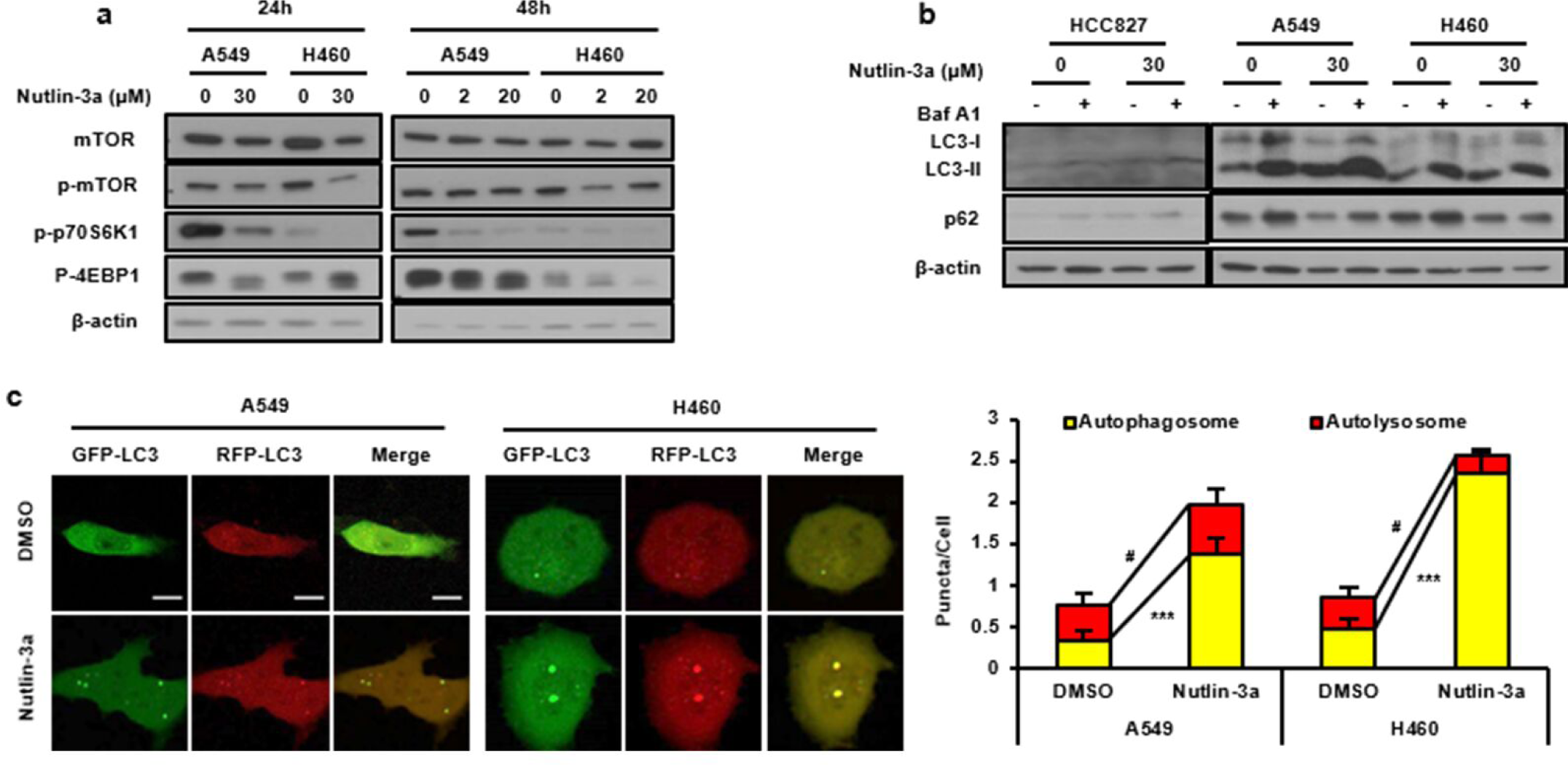
Nutlin-3a disrupts the fusion of autophagosomes and lysosomes in *KRAS* mutant NSCLC cells. **A** Downregulation of the mTOR pathway detected by western blotting after nutlin-3a treatment. **B** Western blotting for the expression of autophagy markers performed after co-treatment of cells with bafilomycin A1 (100 nM) for 1 h in the presence or absence of nutlin-3a for 24 h. **C** Autophagy flux after cell treatment with nutlin-3a (30 μM) for 24 h after transfection of the mRFP-GFP-LC3 plasmid. Live cell imaging was obtained using a confocal laser scanning microscope (left), and number of puncta per cell were quantified from representative images (right) (n≥21). Scale bar: 10 μm.

### Nutlin-3a induces macropinosome derived huge vacuoles

The examination of live cell morphology via light microscopy and 3D holotomography microscopy revealed formation of massive cytoplasmic vacuolization in *KRAS* mutant NSCLC cells, A549, and H460, but not in *KRAS* wild type cells, after treatment with nutlin-3a (Fig 3A and B; Movie EV1). Time-lapse videos shot using a holotomography microscope after treatment with nutlin-3a displayed a ruffling plasma membrane forming macropinocytic cups, and small vacuoles merging to form huge vacuoles with time, finally leading to the rupture of cells (Fig 3C and Movie EV2). Further analysis of whether the nutlin-3a-induced formation of huge vacuoles in the cytoplasm is associated with macropinosomes, which eventually led to fusion with lysosomes for degradation, was conducted. Live cells were examined under a confocal laser scanning microscope after adding of bulk fluid-phase tracer dextran to the medium and staining with a LysoTracker. Cells treated with nutlin-3a had dextran incorporated into huge vacuoles that were not fused with the LysoTracker (Fig 3D). Also, Rab7 and LAMP1, late endosomal markers, were examined under a confocal laser scanning microscope after staining. Both Rab7 and LAMP1 were recruited around huge vacuoles, indicating that nutlin-3a-induced vacuoles had late endosomal characteristics originating from macropinocytosis (Fig 3E and F). Actin cytoskeleton was examined via immunofluorescence staining of Rac1 and phalloidin, because Rac1 activation is required for macropinosome formation and actin polymerization (i.e., for processes such as the formation of lamellipodia and membrane ruffles). In *KRAS* mutant NSCLC cells, Rac1 and phalloidin aggregated outside the membrane, forming protrusions and ruffling with upregulated Rac1 expression after nutlin-3a treatment (Fig 3G and H). Taken together, treatment with nutlin-3a induced actin cytoskeleton remodeling that led to macropinocytosis in *KRAS* mutant NSCLC cells but not in *KRAS* wild type, HCC827, and BEAS-2B cells (Fig EV1).

**Figure 3.**
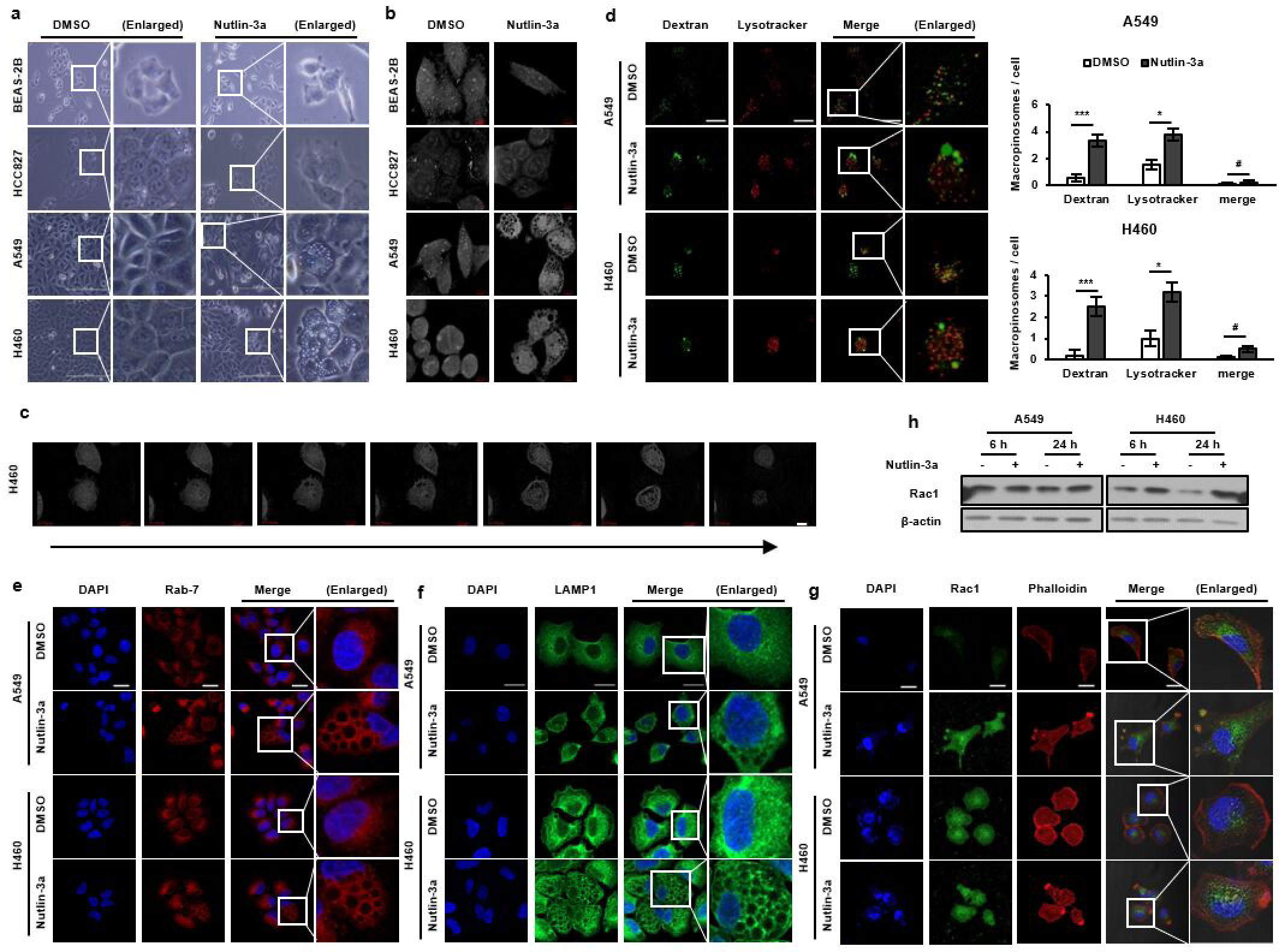
**Nutlin-3a induces huge cytoplasmic vacuoles in *KRAS* mutant NSCLC cells.** Cells were treated with nutlin-3a (30 μM) for 6 (d,h) or 24 h (a,b,e-g). **A, B** Representative live cell images captured by light (a) and holotomography microscope (b) show huge cytoplasmic vacuoles in *KRAS* mutant but not wild type cells. **C** Time-lapse video capture images (24 h) of H460 obtained by holotomography microscopy show increase in macropinosomes volume with time and membrane rupturing in the end. **D** Nutlin-3a induced formation of huge vacuoles were incorporated with dextran (green), but not merged with LysoTracker (red). Representative live images were obtained by confocal laser scanning microscopy (left). Number of macropinosomes were quantitated from the captured image (right) (n≥8). **E-G** Representative images obtained by confocal laser scanning microscopy. Fixed cells were stained with Rab7 (e), LAMP1 (f), or Rac1 (green), and rhodamine–phalloidin (red) (g) and were counterstained with DAPI (blue). **H** Western blotting of Rac1 expression. *p<0.05, **p<0.01, ***p<0.001 compared with control. Scale bar: 200 μm (a), 7 μm (b–c), and 20 μm (d–g).

### Nutlin-3a induces methuosis-like cell death

To investigate whether nutlin-3a induces *KRAS* mutant NSCLC cell death, the levels of apoptosis markers, including cleaved forms of caspase-9, caspase-3, and PARP, were examined using western blotting (Fig 4A). No evident cleaved forms of caspase-9, caspase-3, or PARP were observed after nutlin-3a treatment, compared with that after etoposide treatment, which was used as an apoptosis inducer. In addition, co-treatment with Z-VAD (OMe)-FMK, a pan-caspase inhibitor, did not rescue the loss of cell viability or formation of huge vacuoles induced by nutlin-3a treatment, indicating that nutlin-3a-induced cell death was caspase-independent and non-apoptotic (Fig 4B). To further determine whether nutlin-3a induced cell death, cells were co-treated with ferrostatin-1 (a ferroptosis inhibitor) or necrostatin-1 (a necroptosis inhibitor) and nutlin-3a for 24 h. Neither cell vacuolization nor viability were prevented or rescued (Fig 4C, D and EV2). These results confirmed that the cell death caused by nutlin-3a could not be categorized as apoptosis, ferroptosis, or necroptosis. Furthermore, cells were co-treated with previously known or used macropinocytosis inhibitors, EIPA (Na^+^/H^+^ exchanger inhibitor), which suppress Rac1 signaling by lowering submembranous pH, resulting in the blocking of macropinocytosis; EHT1864 (Rac family small GTPase inhibitor); and bafilomycin A1 (Vacuolar H^+^ ATPase inhibitor) along with nutlin-3a, to determine whether cell death could be rescued. Co-treatment of cells with EIPA, EHT1864, or bafilomycin A1 and nutlin-3a prevented vacuolization but did not rescue viability (Fig 4E-I and Fig EV2). Therefore, it was concluded that nutlin-3a induced macropinocytosis derived methuosis-like cell death, likely due to unknown novel mechanisms.

**Figure 4.**
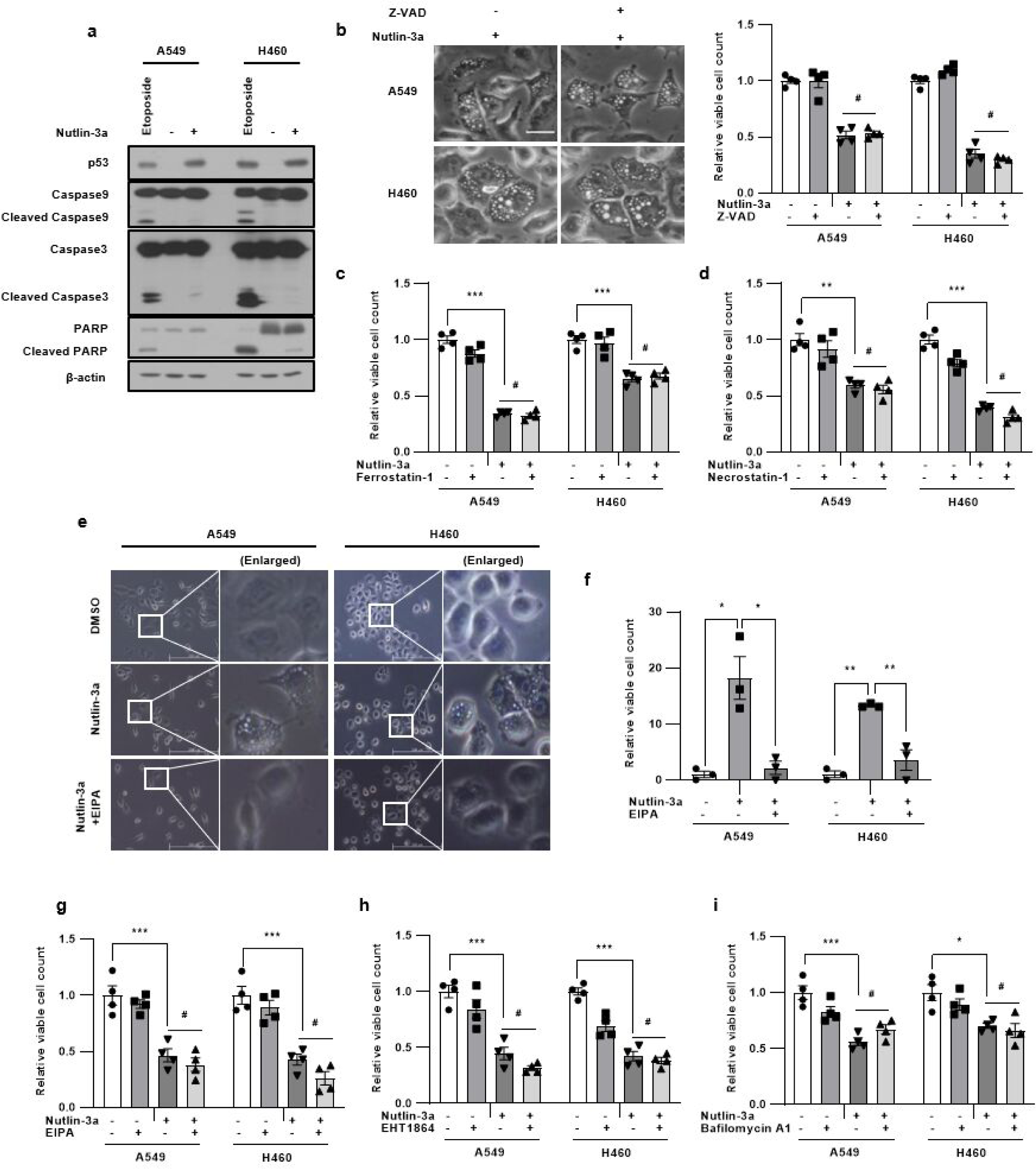
**Nutlin-3a induces methuosis-like death of *KRAS* mutant NSCLC cells.** Cells were treated with nutlin-3a (30 μM) with or without indicated inhibitors for 24 h. **A** Cleavage forms of caspase-9, caspase-3, and PARP did not detected by western blotting. Etoposide treatment (100 μM) for 24 h was used as a positive control for inducing caspase cleavage. **B-I** Representative light microscope images are shown. Cell proliferation was determined via cell counting assay. Cells were co-treated with Z-VAD (OMe)-FMK (50 μM) (b), ferrostatin-1 (1 μM) (c), necrostatin-1 (10 μM) (d), or EHT1864 (10 μM) (h) or pretreated with EIPA (5 μM) for 1 h (e-g) or bafilomycin A1 (5 nM) for 2 h (i) before nutlin-3a treatment. Scale bar: 100 μm for b, and 200 μm for e. *p<0.05, **p<0.01, ***p<0.001 compared with control.

### Nutlin-3a regulates the HBP via GFPT2 suppression

RNA-sequencing (RNA-seq) was performed to identify the nutlin-3a target marker(s) and/or pathways specific to *KRAS* mutant NSCLC cells. Differentially expressed gene sets (5,856 genes) were analyzed between *KRAS* mutant and wild type NSCLC cells and BEAS-2B cells. Venn diagrams were generated to display lists of genes that were upregulated and downregulated in *KRAS* mutant NSCLC compared to those in *KRAS* wild type cells upon nutlin-3a treatment (Fig EV3; Tables EV2 and EV3). Among these genes, the expression of *GFPT2,* glutamine-fructose-6-phosphate transaminase 2, was significantly downregulated only in *KRAS* mutant NSCLC cells, which was confirmed via RT-qPCR (Fig 5A). GFPT2 regulates the flux of glucose into the HBP, which is involved in the UDP-N-acetylglucosamine (UDP-GlcNAc) metabolic process, regulating the availability of precursors for N- and O-linked glycosylation of proteins as well as for O-GlcNAcylation (O-GlcNAc). The level of HBP metabolites were tested via UHPLC–MS/MS, which showed that the abundance of HBP metabolites significantly decreased after treatment with nutlin-3a (Fig 5B). The O-GlcNAc, and N-glycosylation, which use UDP-GlcNAc as a substrate, were examined using western blotting, which revealed that the levels were significantly reduced after nutlin-3a treatment in *KRAS* mutant NSCLC cells (Fig 5C). These results indicate that nutlin-3a regulates the HBP via GFPT2 in *KRAS* mutant NSCLC cells.

**Figure 5.**
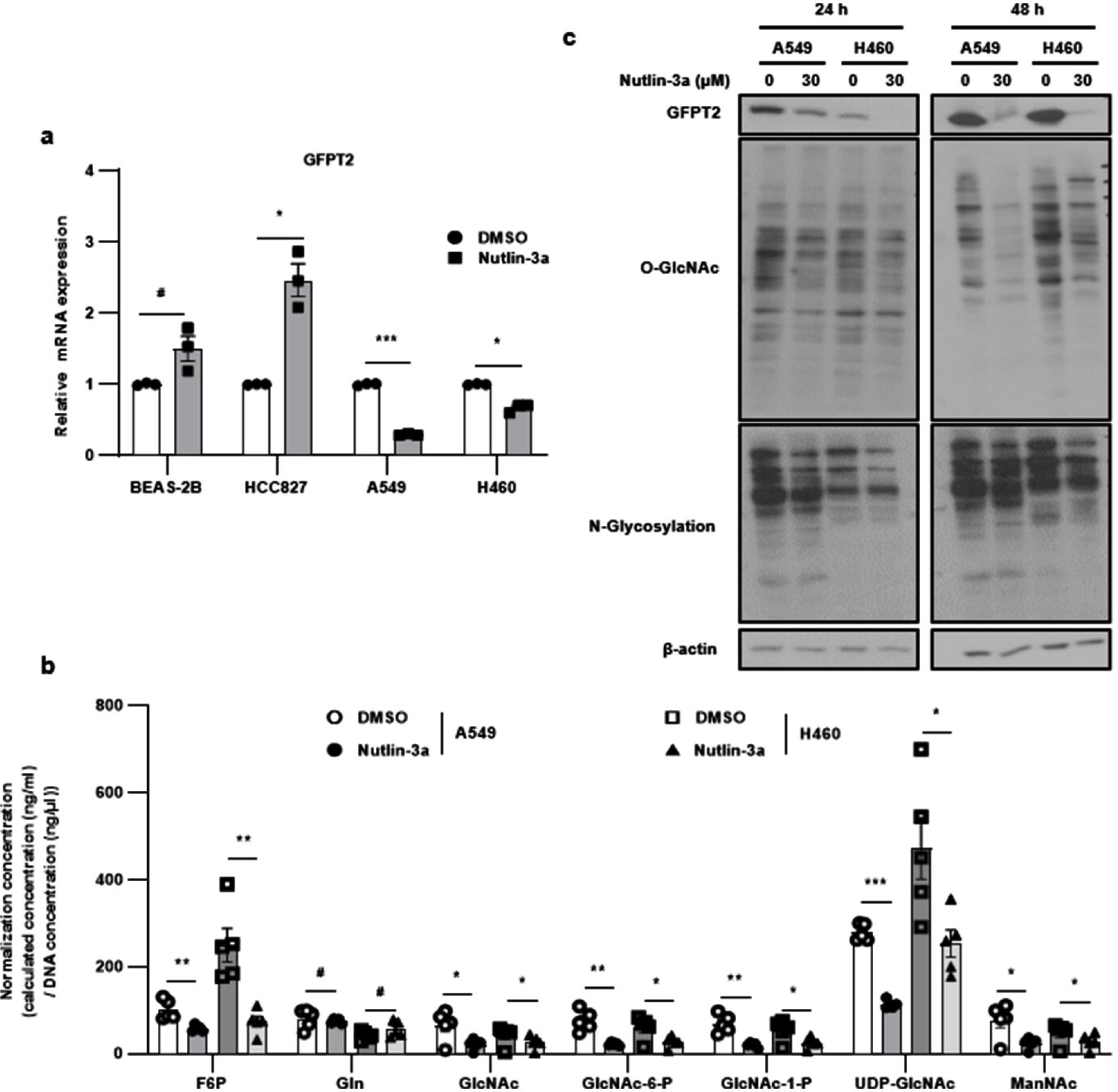
**Downregulation of GFPT2 expression by nutlin-3a alters amounts of HBP metabolites in *KRAS* mutant NSCLC cells. a,b,** Cells were treated with nutlin-3a (30 μM) for 24 h. Expression of *GFPT2* mRNA determined by RT-qPCR **A** Abundance of HBP metabolites analyzed by UHPLC–MS/MS **B,C** Global O-GlcNAcylation of proteins and N-glycosylation with the terminal glycosyl/mannosyl binding lectin examined via western blotting. *p<0.05, **p<0.01, ***p<0.001 compared with control.

### Nutlin-3a-induced methuosis-like cell death is related to the HBP

As treatment with nutlin-3a reduced the amounts of the HBP’s downstream metabolites via GFPT2 suppression, it was assessed if supplementing the downstream HBP metabolites GlcNAc and GlcN rescue nutlin-3a induced cell death via cell counting and annexin V/PI staining. Supplementation with GlcNAc as well as GlcN rescued cell proliferation and death induced by nutlin-3a (Fig 6A and B), suggesting that the HBP is required for *KRAS* mutant NSCLC cell viability. Examination of cell morphology via holotomography microscopy also revealed no vacuole formation or accumulation (Fig 6C and Movie EV3). Furthermore, dextran uptake and LysoTracker staining were assessed to determine if supplementation with GlcNAc reduced macropinosome formation induced by nutlin-3a. GlcNAc supplementation reduced dextran uptake, as observed in controls (Fig 6D). Autophagic flux disruption induced by nutin-3a was rescued by GlcNAc supplementation, showing attenuated yellow puncta formation (Fig 6E). These results indicate that the HBP is associated to macropinosome formation and the fusion of macropinosomes and autophagosomes with lysosomes.

**Figure 6.**
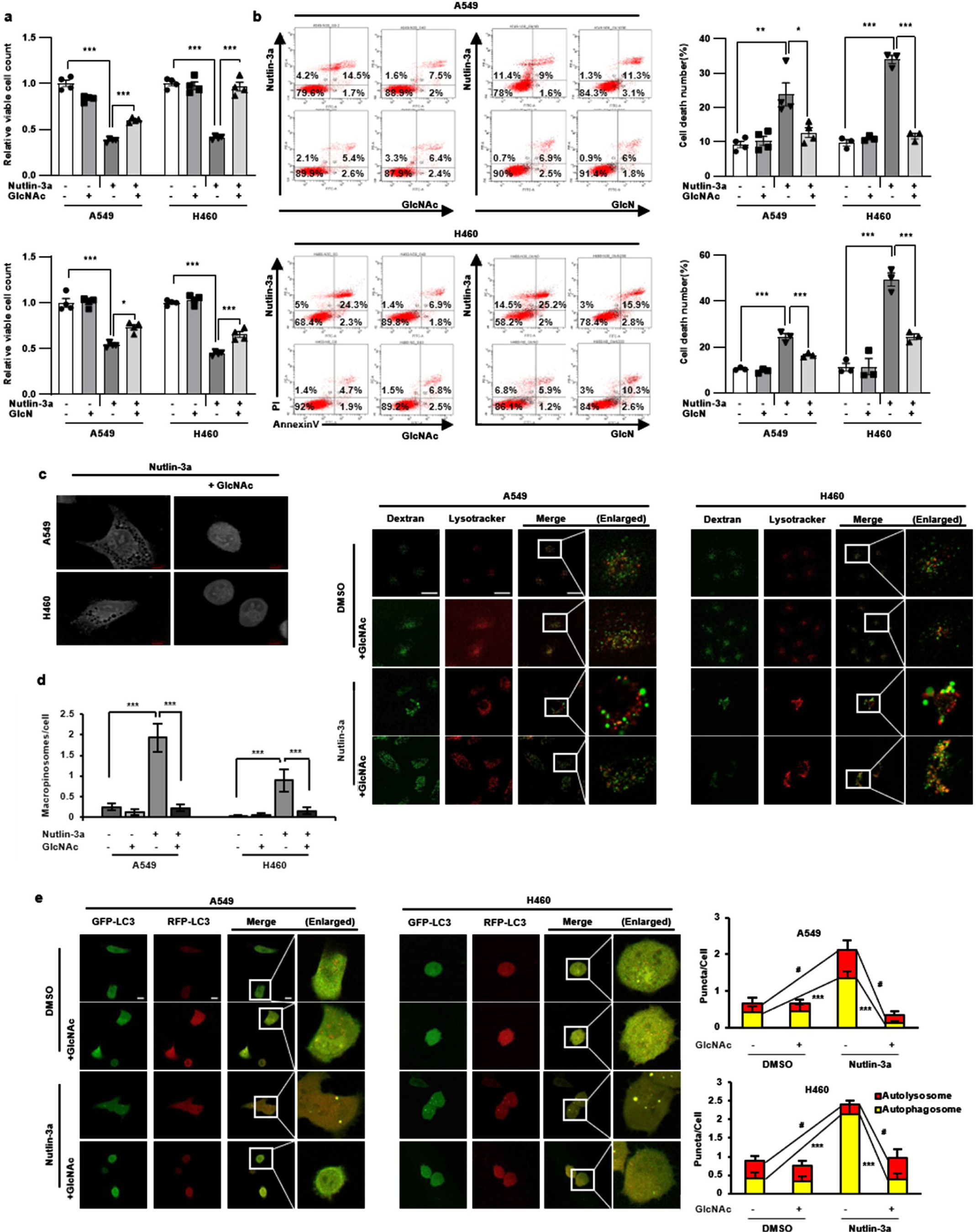
**Nutlin-3a-induced methuosis-like cell death is associated with the HBP in *KRAS* mutant NSCLC cells.** Cells were supplemented with GlcNAc (40 mM) or GlcN (200 μM) for 24 h with or without nutlin-3a (30 μM). **A, B** Both GlcNac and GlcN rescue cell proliferation and death, which was analyzed by cell counting (a), and flow cytometry using annexin V/PI staining (b). **C** Live cell images captured by holotomography microscopy show that GlcNAc supplementation rescues the formation of huge cytoplasmic vacuoles induced by nutlin-3a. **D** Cells were incubated with dextran and LysoTracker for 2 h with GlcNAc or nutlin-3a. GlcNAc supplementation with nutlin-3a recovers dextran uptakes compared to only nutlin-3a treated cells (n≥22). **E** Autophagy flux disruption recovered by supplementation of GlcNAc with nutlin-3a, which was detected after transfection of mRFP-GFP-LC3 tandem vector (n≥20). **D, E** Representative images obtained by confocal laser scanning microscopy and quantitated from the captured image. Scale bar: 7 μm for **c**, and 10 μm for **d**–**e**. *p<0.05, **p<0.01, ***p<0.001 compared with control.

### GFPT2 is a target of nutlin-3a induced cell death

*GFPT2* was knocked down by exogenously introducing siGFPT2, to verify that the HBP enzyme GFPT2 was a target of nutlin-3a in *KRAS* mutant NSCLC cells. Inhibition of O-GlcNAcylation, the KRAS-PI3K/AKT-mTOR pathway (Fig 7A and B) and KRAS-GTP activity (Fig 7C) were verified using western blotting. Cell proliferation and viability were determined using MTT assay (Fig 7D) and annexin V/PI staining (Fig 7E). GFPT2 inhibition suppressed both proliferation and viability in the manner of caspase-independent cell death mechanism, as shown after nutlin-3a treatment (Fig 7F). These results indicate that GFPT2 of the HBP, which is a target of nutlin-3a, plays an important role in the viability of KRAS mutant NSCLC cells.

**Figure 7.**
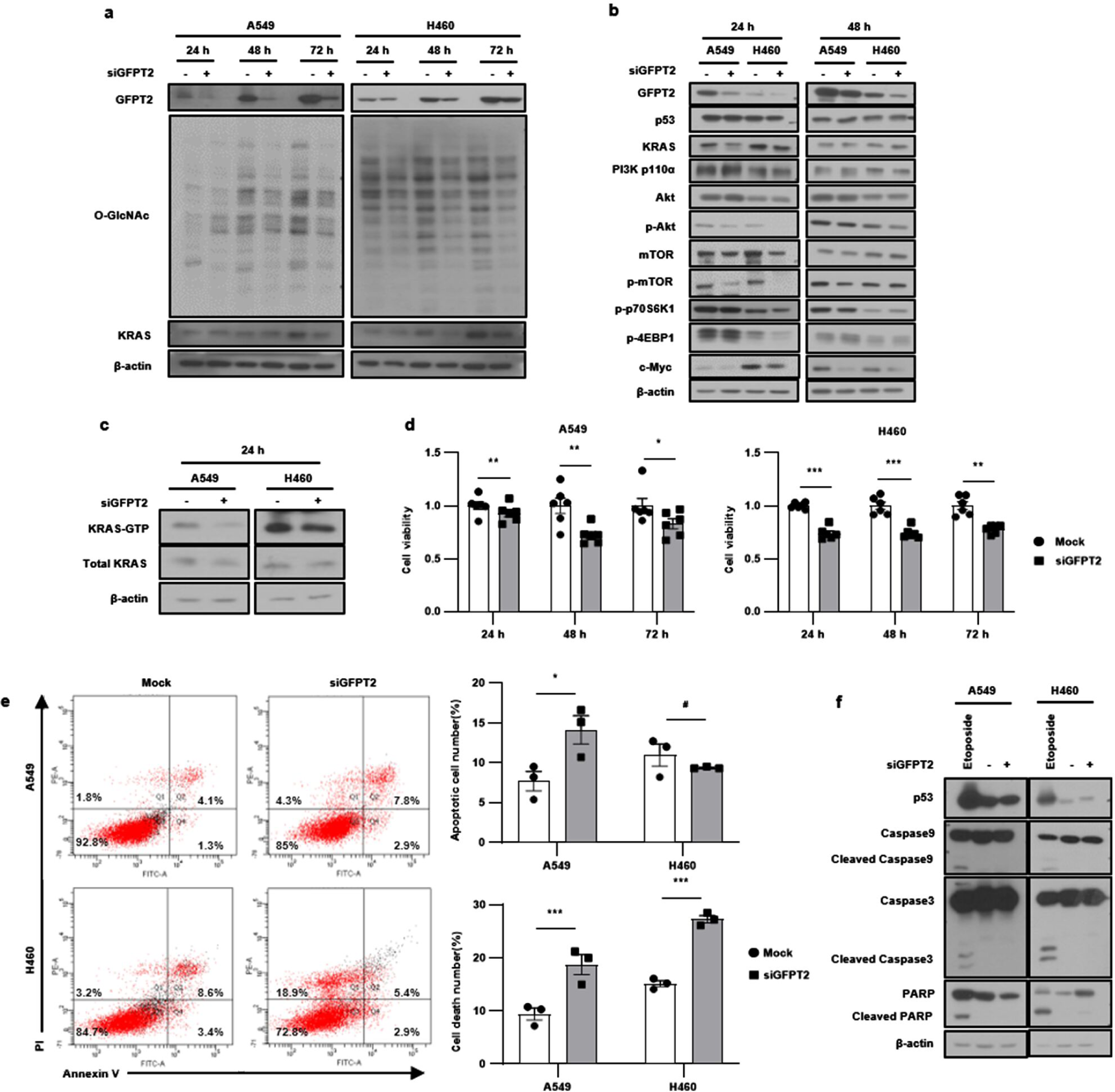
GFPT2 is a target of nutlin-3a-induced cell death. GFPT2 was inhibited by exogenously introduced siGFPT2 for indicated times in *KRAS* mutant NSCLC cells. **A, B** Expression of GFPT2 (a), O-GlcNAc and, the KRAS-PI3K/Akt-mTOR pathway (b) detected by western blotting. **C** KRAS-GTP form detected through the KRAS activation assay. **D** Cell viability measured by MTT assay. **E** Apoptotic cell death number (top), total cell death number (bottom) analyzed via flow cytometry using annexin V/PI staining. **F** Expression of caspase-9, caspase-3, and PARP detected by western blotting. *p<0.05, **p<0.01, ***p<0.001 compared with control.

### Nutlin-3a inhibits tumor growth *in vivo*

Xenograft zebrafish and mouse models were used to investigate anticancer effect of nultin-3a *in vivo*. CM-Dil-labeled (red) cells were transplanted into zebrafish embryos and cultured with DMSO, nutlin-3a, and/or GlcNAc supplemented with nutlin-3a for 5 days. Nutlin-3a treatment reduced the tumor area of both cell types, whereas GlcNAc supplementation rescued this effect (Fig 8A). Consistently, nutlin-3a reduced tumor growth (Fig 8B, C and EV4A–C) in mouse xenografts without any difference in body weight (Fig 8D and EV 4D). After final treatment with nutlin-3a, mice were euthanized, and tumor tissues were subjected to western blotting and immunohistochemical analyses. GFPT2 expression was downregulated and p53 expression was upregulated in nutlin-3a-treated tumor tissues (Fig 8G, EV4E and H). Tumor proliferation, determined by Ki67 staining, was also decreased after nutlin-3a treatment (Fig 8E and EV4F). In addition, vacuoles from the H&E stained nutlin-3a-treated tumor tissue showed the formation of large vacuoles compared to their control counterparts (Fig 8F and EV4G), confirming the anticancer effects of nutlin-3a *in vivo*.

**Figure 8.**
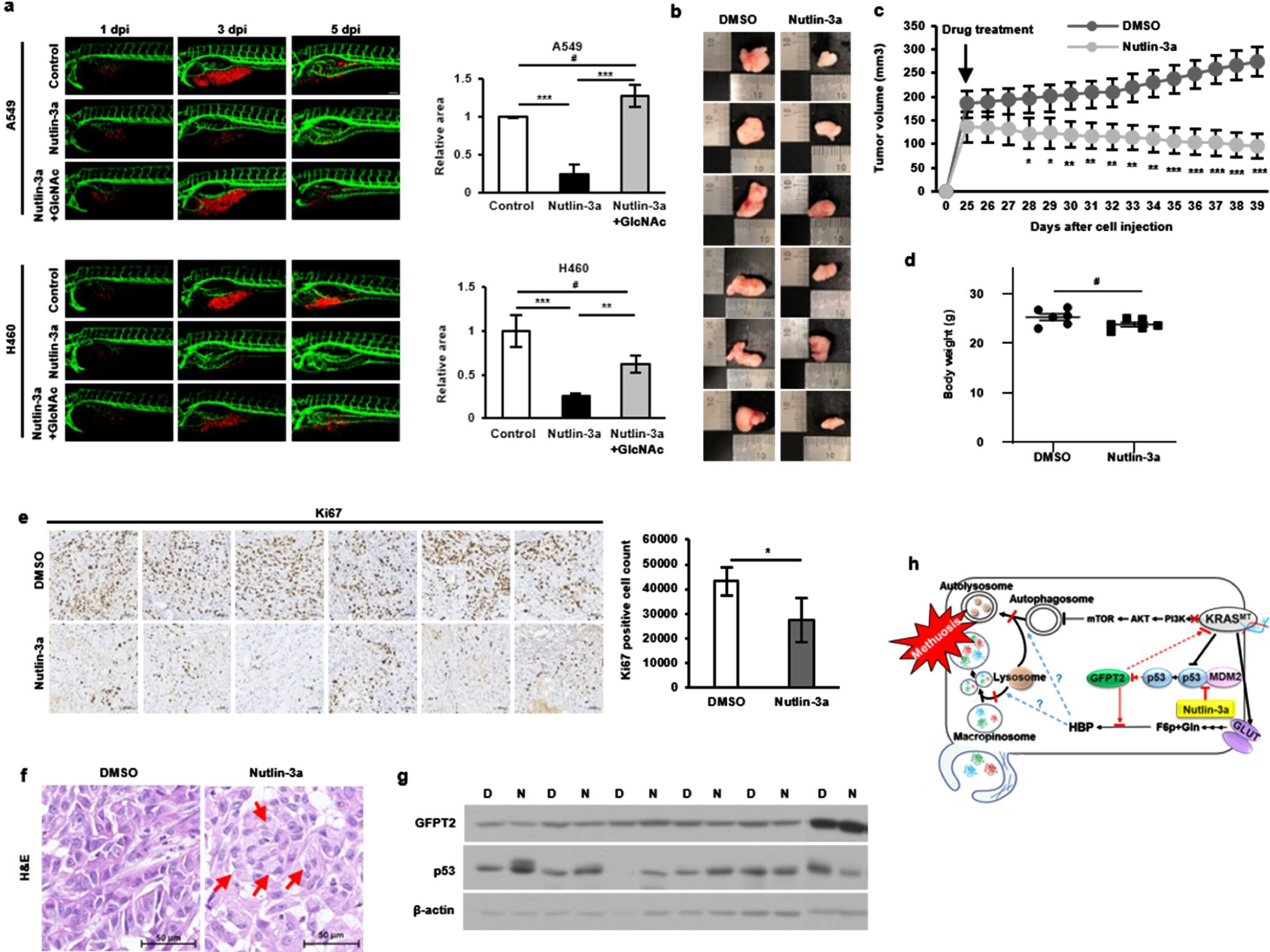
Anticancer effects of nutlin-3a confirmed *in vivo*. **A** Representative confocal image of CM-Dil-labeled cancer cells (red) and vasculature (green) in zebrafish larvae (left). Quantified data show cancer cell volume after nutlin-3a treatment and GlcNAc co-treatment compared with that of the control group (right). Supplement of GlcNAc-rescued cancer cell volume reduces after nutlin-3a treatment. **B, C** Images (b) and comparison (c) of growth rates of subcutaneous tumors formed by A549 injection; tumor growth is observed in the presence or absence of nutlin-3a. **D** Weight of mice after treatment with either vehicle control or nutlin-3a on the day of euthanasia. **E** Immunohistochemical analysis of Ki67 expression in tumor tissues. **F** Vacuole examination in H&E stained tumor tissues. Red arrows indicate vacuoles. **G** Western blot analysis of GFPT2 and p53 expression in *in vivo* xenograft tumors. **H** Proposed mechanism of nutlin-3a-induced methuosis-like cell death. Scale bar: 20 μm for (a) and 50 μm for (e–f). Data are presented as mean ± standard deviation; n = 6. *p<0.05, **p<0.01, ***p<0.001 compared with control.

## Discussion

Nutlin-3a is a small-molecule, MDM2 antagonist that effectively disrupts p53-MDM2 interaction, leading to stabilization and activation of the p53 pathway in p53 wild-type cells, selectively inducing senescence, cell cycle arrest, and apoptosis in various wild-type p53 cancer cells (Vassilev *et al*, 2004; Tovar *et al*, 2006). Some of its derivatives, such as APG-115, BI-907828, HDM201, and milademetan (DS-3032b), are undergoing clinical trials (Yi *et al*, 2018; Hao *et al*, 2022; Stein *et al*, 2022, Takahashi *et al*, 2021). However, the number of studies on the effect of nutlin-3a in lung cancer cells is scarce; most studies show combinational effects with radio- or chemo-therapy (Shen & Maki, 2011; Luo *et al*, 2013; Kocik *et al*, 2019). Therefore, we examined the effect of nutlin-3a on NSCLC cells and discovered that nutlin-3a induces *KRAS* mutant NSCLC cells specific caspase independent death, presenting massive macropinocytosis derived methuosis-like phenotype. No-label live holotomography cell image and time-lapse video by tomoholography microscopy revealed an accumulation of huge vacuoles in the cytoplasm, where small vesicles formed by macropinocytosis merged to form huge vacuoles with time, and then the cell ruptured finally after nutlin-3a treatment. Massive macropinosomes induced by nutlin-3a uptake dextran and were positive for late endosome markers LAMP1 and Rab7; however, they neither shrank nor fused with lysosomes, indicating disruption of the canonical endocytotic pathway along with the alteration of cytoskeleton organization, which involved actin-mediated membrane ruffling. Consistent with these findings, the disruption of the fusion of autophagosomes and lysosomes was evaluated, even though autophagy was increased by nutlin-3a. This indicated that dysfunction of lysosomes and/or alteration of the cytoskeleton may play a role in the methuosis-like cell death induced by nutlin-3a, the detailed mechanism of which needs to be further investigated (Yang *et al*, 2020; Ramirez *et al*, 2019; Vats & Manjithaya, 2019). Co-treatment of nutlin-3a with Z-VAD, ferrostatin-1, and necrostatin validated that nutlin-3a-induced methuosis-like cell death was caspase-independent, non-apoptotic, non-ferroptotic, and non-necroptotic. Contrary to our expectations, co-treatment of nutlin-3a with various known macropinocytosis or methuosis inhibitors, such as EIPA (Koivusalo *et al*, 2010), EHT1864 (Mao *et al*, 2017), and bafilomycin A1 (Harada *et al*, 1996; Compton *et al*, 2016), did not rescue cell viability. Based on these results, vacuole formation can be rescued by inhibiting membrane ruffle formation or V-ATPase activity. However, there is a possibility that another underlying mechanism may contribute to inducing cell death or that induced macropinocytosis itself may not be the reason for cell death. The majority of previous studies did not investigate the rescue of cell viability, or if tested by treating with the above inhibitors, cell viability was not rescued (Overmeyer *et al*, 2011; Brack *et al*, 2020; Nara *et al*, 2010). The underlying mechanism or pathway of methuosis or methuosis-like cell death remains unclear, but the involvement of Ras (Bhanot *et al*, 2010), Rac1 (Bhanot *et al*, 2010; Chi *et al*, 1999), Akt-mTOR (Liu *et al*, 2021), and AMPK (Bielsa *et al*, 2022) in a caspase-independent manner has been reported. Consistent with these reports, nutlin-3a downregulated KRAS activity as well as the KRAS downstream signal, KRAS-PI3K/Akt-mTOR, but not MEK/ERK. Nutlin-3a decreased levels of B-Raf and C-Raf, but not those of p-MEK and p-ERK, the reasons for which have not yet been elucidated. However, it may have bypassed the MEK/ERK pathway activation by downregulation of the PI3K/Akt pathway to maintain the balance of these two pathways to perform appropriate cellular function (Hayashi *et al*, 2008).

To further understand the underlying mechanism and identify the target of nutlin-3a function, RNA-seq was performed. GFPT2 were found to be downregulated in *KRAS* mutant NSCLC cells. The HBP, which is regulated by GFPT2, is a UDP-GlcNAc metabolic process, and its important end product, UDP-GlcNAc, is required for the O-GlcNAcylation and N-and O-linked glycosylation of proteins (Helenius & Aebi, 2001; Biwi *et al*, 2018). Following examination showed that amounts of O-GlcNAc, N-glycosylation, and HBP-related metabolites, fructose 6-phosphate (F6P), glucosamine (Gln), GlcNAc, GlcNAc-6-phosphate (GlcNAc-6-P), GlcNAc-1-phosphate (GlcNAc-1-P), UDP-GlcNAc, and N-acetylmannosamine (ManNAc) were reduced along with GFPT2 suppression by nutlin-3a. In addition, supplementation with the HBP metabolites such as GlcNAc or GlcN rescued the nutlin-3a induced methuosis-like cell death as well as disruption of autophagic flux in KRAS mutant NSCLC cells. These results suggest that GFPT2 is a crucial component in the HBP pathway and a key player in the process of methuosis-like cell death induced by nutlin-3a. This was validated by experiments with the knockdown of GFPT2. Suppression of GFPT2 may induce massive macropinocytosis due to the blocking of the HBP, which leads to a lack of the necessary metabolites for cell survival, especially in *KRAS* mutant cancer cells because macropinocytosis is an important process for nutrient uptake (Recouvreux & Commisso, 2017). KRAS mutant NSCLC specific cell death induced by nutlin-3a, confirmed *in vivo* in zebrafish and mouse models and rescued by GlcNAC supplementation, was verified using *in vivo* zebrafish xenograft models.

GFPT2 (GFAT2), which has an amino acid homology of 75.6% with GFPT1 (GFAT1), is a rate-limiting enzyme in the HBP, similar to GFPT1, and is responsible for glycosylation. Recently, the role of GFPT2 in tumorigenesis, such as its association with aggressiveness and EMT/migration, has been gradually recognized in various cancers, including breast, colon, and lung cancer (Taparra *et al*, 2018; Simpson *et al*, 2012; Liu *et al*, 2020). Along with GFPT2, various functions of the HBP as well as the O-GlcNAc, which is a post-translational modification using UDP-GlcNAc, the end product of HBP, as a substrate, have also been proposed as targets for cancer treatment (Taparra *et al*, 2018; Ricciardiello *et al*, 2018; Chen *et al*, 2019; Kim *et al*, 2020). In particular, oncogenic KRAS mutations drive glucose uptake and utilization via metabolic reprogramming to support cancer cell growth, which enhances the flux of glucose to the HBP and glycolytic biosynthetic branches (Ying *et al*, 2012) ^49^. Thus, it is advisable to consider the role of aberrant HBP through the dysfunction of its enzyme, resulting in abnormal O-GlcNAc in *KRAS* mutant cancer, as supported by recent studies (Taparra *et al*, 2018; Kim *et al*, 2020). Increased or decreased O-GlcNAcylation, which stabilizes proteins such as snail1, β-catenin, and c-Myc (Park *et al*, 2010; Olivier-Van Stichelen *et al*, 2014; Chou *et al*, 1995), by suppression of GFPT2 of the HBP, showed delayed tumorigenesis in KRAS-MT lung cancer (Kim *et al*, 2020). Consistent with this, decreased c-Myc expression along with the reduction of O-GlcNAc by nutlin-3a was observed in our results; the underlying mechanism and function during tumorigenesis remain unclear and need to be evaluated.

Overall, the therapeutic efficacy of *KRAS mutant* NSCLC is poor. However, direct *KRAS*-G12C inhibitors, such as sotorasib, and MRTX849 are breakthrough drugs that ensure targeted therapies in patients with *KRAS*-G12C mutant NSCLC. However, the prognosis of patients remains disappointing owing to disease heterogeneity and acquired resistance, leaving room for the development of more effective treatment options. Immunotherapy based on immune checkpoint inhibitors (ICIs) has been one of the options that has shown success in treating certain groups of patients with NSCLC. Although the influence of *KRAS* mutation status on the immune response of patients with NSCLC remains controversial, many studies have suggested that there may be a synergistic effect between *KRAS*-G12C inhibitors and immunotherapy drugs combined with ICIs (Canon *et al*, 2019). Moreover, the impact of post-translational modifications such as N-glycosylation of PD-1 or PD-L1, which is regulated by HBP, on cancer therapy and clinical diagnosis has been suggested (Sharma *et al*, 2020; Chen *et al*, 2021). Therefore, nutlin-3a, which inhibits the HBP via GFPT2, may be an effective therapeutic option in combination with ICIs, for the treatment of NSCLC.

In conclusion, nutlin-3a induced the downregulation of the KRAS/PI3K/mTOR pathway and methuosis-like cell death by disrupting the fusion of lysosomes with both macropinosomes and autophagosomes, which is reliant on the suppression of GFPT2 in the HBP of KRAS mutant NSCLC cells (Fig. 8h). This suggests that nultin-3a has a significant potential for treating KRAS mutant NSCLC cells and overcoming the resistance developed by cancer cells, which currently has limited treatment options, even though the underlying mechanism needs to be further investigated.

## Materials and Methods

### Cell culture and reagents

One normal human bronchial epithelial cell line (BEAS-2B) and ten NSCLC cell lines (HCC827, H1299, H1975, PC9, PC9GR, H820, A427, A549, H460, and H358) were purchased from the American Type Culture Collection (Manassas, VA, USA) and the Korean Cell Line Bank (Seoul, South Korea). All cells, except for A427, which was cultured in Dulbecco’s modified Eagle’s medium (Hyclone, Logan, UT, USA), were cultured in RPMI 1640 medium (WELGENE, Daegu, South Korea) supplemented with 10% fetal bovine serum (WELGENE), and 1% penicillin/streptomycin (WELGENE) at 37℃ under 5% CO₂ conditions. Nutlin-3a (SML0580; Sigma–Aldrich, St. Louis, MI, USA), etoposide (S1225, Selleckchem, Houston, TX, USA), Z-VAD-FMK (S7023, Selleckchem), bafilomycin A1 (B1793, Sigma–Aldrich), EIPA (14406, Cayman Chemical Company, Michigan, USA), EHT1864 (S7482, Selleckchem), and LY294002 (L9908, Sigma–Aldrich) were purchased and dissolved in DMSO (D2650, Sigma-Aldrich). Endoribonuclease-prepared small interfering RNAs (esiRNAs; EHU144601; Sigma-Aldrich) were used for *GFPT2* knockdown.

### Cell proliferation and viability

Cells (3-5 × 10^5^) were seeded into 6-well plates, and their proliferation was determined via cell counting. After 24 h of culture, the cells were trypsinized and counted using a hemocytometer. Representative images were captured using an inverted light microscope (Olympus IX71, Olympus Corp. Japan). Cell viability was measured using MTT assay. Briefly, cells (1×10^3^) were seeded into 96-well plates and treated with nutlin-3a for 24 h. Then, 50 μl of MTT solution (0.5 mg/ml, Sigma–Aldrich) was added to each well, and the plate was incubated for 2 h. The MTT solution was removed, and 100 μl of DMSO (Samchun chemicals, Seoul, South Korea) was added to each well. The absorbance was measured at 570 nm on a microplate reader (SpectraMax Plus 384, Molecular Devices, USA).

### Colony formation assay

Cells (500) were seeded into 6-well plates. After 14 days of culture, visible colonies were fixed and then stained with Hemacolor ® rapid staining solutions (Merck & Co., Inc., Rahway, NJ, USA) for 5 min. Images were acquired using a scanner (SCX8123, Samsung, Suwon, South Korea), and colony numbers were counted using ImageJ (version 1/52n, NIH, Bethesda, MD, USA).

### Flow cytometry analysis

Cells (5 × 10^5^) were seeded into 6-cm dishes and incubated for 24 h. For cell cycle analysis, nutlin-3a-treated cells were fixed with ice-cold 70% ethanol, incubated at −20℃ for at least 1 h, washed with PBS and then resuspended in 2 mM EDTA buffer in PBS with 0.1 mg/mL RNase A and 50 mg/mL PI (P4864, Sigma-Aldrich) for 20 min at ∼ 20℃ in the dark. Stained samples were subjected to flow cytometry analysis (BD FACSCanto™ II, Becton, Dickinson and Company, USA) within 1 h. To examine cell death, cells were treated with nutlin-3a in the absence or presence of GlcNAc and GlcN or transfected with siGFPT2 for 24 h. The samples were stained using the FITC Annexin V Apoptosis Detection Kit I (BD Pharmingen™, NJ, USA).

### RNA isolation and reverse transcription-quantitative polymerase chain reaction (RT-qPCR)

Total RNA was isolated from cells using TRIzol reagent (Invitrogen, Carlsbad, CA, USA) and reverse-transcribed to cDNA using a cDNA synthesis kit (Cosmo Genetech, Seoul, South Korea), according to the manufacturers’ instructions. RT-qPCR was conducted using gene-specific primers with SYBR Green Q Master (Cosmo Genetech) on an ABI 7500 Real-Time PCR System (Applied Biosystems, Warrington, UK). The primers used were as follows: *GFPT2* forward: 5′ – AGGTGCATTCGCGCTGGTT – 3′, reverse: 5′ – TGTGGAGAGCTTGTATTTGCTCC -3′; *GAPDH* forward: 5′ – ACCCACTCCTCCACCTTTGA -3′, reverse: 5′-CTGTTGCTGTAGCCAAATTCGT -3′. The 2^−ΔΔCT^ values of the target genes were normalized to those of the endogenous control, *GAPDH*.

### Western blot analysis

Proteins from cell lysates were separated via SDS-PAGE and transferred to Immobilon-P PVDF membranes (IPVH00010, Merck & Co., Inc., USA). These membranes were subsequently probed with the indicated primary antibodies and incubated with appropriate goat anti-rabbit IgG or goat anti-mouse IgG (Cell Signaling Technology, USA) secondary antibodies conjugated with horseradish peroxidase (HRP) before signal detection using an enhanced chemiluminescence (ECL) system (Translab, Daejeon, South Korea) according to the manufacturer’s instructions. Primary antibodies against p53 (sc-126), B-Raf (sc-5284), c-Myc (sc-40), O-GlcNAc (sc-59623), and β-actin (sc-47778) were purchased from Santa Cruz Biotechnology (CA, USA). Antibody against KRAS (12063-1-AP) was purchased from Proteintech (Rosemont, IL, USA); those against PI3 kinase p110α (#4249), Akt (#4691), p-Akt (#4060), p-B-Raf (#2696), C-Raf (#9422), p-C-Raf (#9427), MEK 1/2 (#9122), p-MEK 1/2 (#9154), Erk1/2 (#9102), p-Erk1/2 (#9101), caspase-9 (#9502), caspase-3 (#9662), PARP (#9542), mTOR (#2983), p-mTOR (#5536), p-p70S6K1 (#9234), and p-4EBP1 (#2855) were purchased from Cell Signalling Technology (Danvers, MA, USA); those against p62 (ab56416) and GFPT2 (ab190966) were purchased from Abcam (Cambridge, UK); those against Rac1 (610650) were purchased from Becton, Dickinson and Company; and those against LC3 (PB036) were purchased from MBL (Nagoya, Japan).

### N-glycosylation analysis

N-glycosylation was evaluated as described previously ^43^. Briefly, the membrane was incubated at 4°C overnight with concanavalin A–HRP lectin (L6397; Sigma; 0.25 μg/ml in high-salt TBST [1 M NaCl in TBST]) and then washed for 1 h in high-salt TBST, followed by which signal detection was conducted using the ECL system.

### KRAS activation assay

KRAS activation was determined using a commercial KRAS Activation Assay Kit (STA-400-K, Cell Biolabs, USA), according to the manufacturer’s protocol. Briefly, cells were harvested, washed with ice-cold PBS, and lysed in ice-cold 1× assay/lysis buffer containing a 1× protease inhibitor cocktail. The lysates were centrifuged at 14,000 g for 10 min at 4℃. The supernatant (500 μg) was incubated with Raf1 RBD agarose beads to detect KRAS–GTP. The beads were washed three times with 1× assay buffer. The bound proteins were eluted with 2× Laemmli sample buffer and loaded onto an SDS-PAGE gel. The primary anti-KRAS antibody was provided with the kit.

### Immunofluorescence staining

Cells (1 × 10^5^) were seeded onto coverslips placed in a 12-well plate. After 24 h, the cells were washed twice with ice-cold PBS and fixed with 4% formaldehyde for 15 min at RT. The fixed cells were washed thrice with ice-cold PBS for 5 min, after which, the cells were permeabilized with 0.1% TritonX-100 in PBS for 15 min, blocked with 3% BSA, incubated with anti-Rac1, anti-LAMP1 (sc-20011, Santa Cruz Biotechnology), anti-Rab7 (#9367, Cell Signaling Technology) antibodies overnight at 4℃ in the dark, and then incubated with Alexa Fluor 488- or 546-conjugated secondary antibodies. Rhodamine–phalloidin (R415; Invitrogen, Carlsbad, CA, USA) was used to stain F-actin for 1 h in the dark. Each coverslip was mounted with medium containing the nuclear stain DAPI (Vectashield; H1200, Vector Laboratories, USA). Fluorescence images were obtained using confocal laser scanning microscopy (LSM 780 and LSM800, Carl Zeiss, Oberkochen, Germany) and visualized using an appropriated objective lens.

### Dextran uptake analysis

Cells (2 × 10^5^) were seeded into confocal dishes and incubated overnight. The cells were then incubated with dextran (D22910, Sigma) and LysoTracker (L7528, Thermo Fisher Scientific), according to the manufacturer’s protocol. Fluorescence images were obtained using LSM 780 or LSM800 (Carl Zeiss) and visualized using 100× oil or 40× water immersion objective lenses.

### Live holotomography microscopy

Cells (2 × 10^5^) were seeded into a TomoDish (Tomocube Inc., Daejeon, South Korea) and incubated overnight. The 3D label-free live cell images were obtained with a 3D holotomography microscope (HT-2H, Tomocube Inc.) at 37℃ in a 5% CO_2_ atmosphere at a wavelength of 532 nm. The cells were visualized based on the reflective index measurements, which allowed label-free imaging.

### Autophagic flux measurement

Autophagic flux was detected in live cells using the mRFP-GFP-LC3 tandem vector kindly provided by Professor T. Yoshimori (Department of Genetics, Graduate School of Medicine, Osaka University, Japan). Cells cultured in a 6-cm dish were transfected for 48 h with the mRFP-GFP-LC3 tandem vector. Fluorescence images were obtained using LSM 780 or LSM800 (Carl Zeiss) and visualized using an appropriated objective lens.

### RNA-sequencing

Total RNA was extracted from samples using TRIzol reagent (Invitrogen), according to the manufacturer’s protocol. Total mRNA-seq was performed by Macrogen (Seoul, South Korea).

### Metabolomics

Cells (2–4 × 10^6^) were seeded into a 10-cm plate, incubated overnight, and then treated with nutlin-3a for 24 h. For cellular metabolite extraction, ice-cold 50% methanol containing an internal standard (glutamine-d5, 1 μg/mL) was used. The cell pellets were lysed via freeze/thaw using liquid nitrogen, and the lysate was centrifuged at 14,000 rpm at 4°C for 10 min. The supernatant was then reconstituted and injected into the UPLC–MS/MS system. Quantitative analysis was conducted using an LC–MS/MS system consisting of an AB Sciex Exion LC Series UHPLC (AB Sciex, Redwood City, CA, USA) and an AB Sciex 4500 Triple Quad mass spectrometer (AB Sciex). Chromatographic separation of HBP metabolites was achieved using a hydrophilic interaction liquid chromatography (HILIC) column, and the mobile phase was eluted by gradient elution. To detect metabolites, an AB Sciex 4500 Triple Quad mass spectrometer equipped with an electrospray ionization source in the negative ionization mode was used. All analytes were detected by multiple reaction monitoring, and all data processing was performed using the Analyst software (version 1.6.2, AB Sciex).

### *In vivo* zebrafish tumor model

Zebrafish (*Danio rerio*) experimental protocols were approved by the local ethics board (Sookmyung Women’s University Animal Care and Use Committee, SMWU-IACUC-1712-036) and performed as previously described (Park *et al*, 2018). One day post-injection (dpi), the embryos were administered 7 μM nutlin-3a or co-treated with GlcNAc 40 mM for 5 days.

### *In vivo* xenograft mouse model

All experimental procedures and animal care were performed as previously described (Park *et al*, 2018) in accordance with the protocols approved by the Institutional Animal Care & Use Committee, Korea University (IACUC no. KOREA-2021-0005). Briefly, cells (1 × 10^7^) were subcutaneously injected into the flanks of 7-week-old BALB/c nude mice (Orient Bio, Korea). After cell injection, nutlin-3a was injected intraperitoneally at a concentration of either 25 mg/kg or 10% DMSO in PBS when the xenografted tumors measured approximately 100 mm^3^. Injections were administered every other day for 2 weeks, and the tumor size was measured using an electronic caliper on the same schedule.

### Mouse tumor tissue analysis

Mouse tumor tissues were divided into two fragments and either fixed or frozen in nitrogen. The frozen tissues were homogenized in lysis buffer and used for western blotting. The other tissue fragments were fixed in 10% neutral-buffered formalin and embedded in paraffin. The tissues were stained using H&E or a primary antibody against Ki67 (GA626, Dako Omnis) and an HRP-labeled secondary antibody (Abcam), according to the manufacturers’ protocols. Images were analyzed using Axio Scan Z1 (Carl Zeiss).

### Statistical analysis

Experimental results from at least three independent experiments are presented as mean ± standard deviation. GraphPad Prism 9 software (GraphPad, CA, USA) and IBM SPSS statistics (version 25.0; IBM, NY, USA) were used for statistical analyses. A two-tailed Student’s *t*-test was conducted to compare two independent groups, and ANOVA was used to compare multiple groups. If the p-value obtained by ANOVA was <0.05, the p-values between the groups were compared with post-hoc analyses using Tukey’s HSD. Significance was defined as follows: #0.05<p (no significance), * p<0.05, ** p<0.01, *** p<0.001; p <0.05 was considered to indicate significant difference.

## Acknowledgments

This work was supported by grants from the Basic Science Research Program (NRF-2019R1A2C1083909 and NRF-2023R1A2C1002372) of the National Research Foundation of Korea, which is funded by the Ministry of Education, Science, and Technology of South Korea. We thank the Korea Basic Science Institute (KBSI) imaging team and the Korean Biomachine Facility for technical support in using Tomocube, Inc. KBSI under the R&D programs (C080200) was supervised by the Ministry of Science and ICT, Republic of Korea.

## Author contributions

D.K. and J.L. conceived and designed the experiments; D.K., D.M., and J.K. conducted the experiments; D.K., D.M., J.K., and J.L. analyzed and interpreted the results. J.K. and M.J.K. performed the zebrafish xenograft experiments and analysis. Y.S. and B.H.J. developed the LC–MS/MS methods and performed the analyses. D.K. and J.L. wrote the manuscript, and all authors reviewed the manuscript.

